# Emergence of saliva protein genes in the secretory calcium-binding phosphoprotein (SCPP) locus and accelerated evolution in primates

**DOI:** 10.1101/2024.02.14.580359

**Authors:** Petar Pajic, Luane Landau, Omer Gokcumen, Stefan Ruhl

**Author notes:** **correspondence:** O.G., S.R.

## Abstract

Genes within the secretory calcium-binding phosphoprotein (SCPP) family evolved in conjunction with major evolutionary milestones: the formation of a calcified skeleton in vertebrates, the emergence of tooth enamel in fish, and the introduction of lactation in mammals. The SCPP gene family also contains genes expressed primarily and abundantly in human saliva. Here, we explored the evolution of the saliva-related SCPP genes by harnessing currently available genomic and transcriptomic resources. Our findings provide insights into the expansion and diversification of SCPP genes, notably identifying previously undocumented convergent gene duplications. In primate genomes, we found additional duplication and diversification events that affected genes coding for proteins secreted in saliva. These saliva-related SCPP genes exhibit signatures of positive selection in the primate lineage while the other genes in the same locus remain conserved. We found that regulatory shifts and gene turnover events facilitated the accelerated gain of salivary expression. Collectively, our results position the SCPP gene family as a hotbed of evolutionary innovation, suggesting the potential role of dietary and pathogenic pressures in the adaptive diversification of the saliva composition in primates, including humans.

## Introduction

The secretory calcium-binding phosphoprotein (SCPP) family of genes has been of interest for its regulation of calcium-phosphate concentration in tissues as diverse as bone, dentin, enamel, mammary glands, and salivary glands [1]. The SCPP gene repertoire, comprising more than 20 genes in humans, was proposed to have originated from a single ancestral gene, *SPARCL1*, involved in the mineralization of skeletal tissue in bony fish [2, 3]. SCPP genes evolved as calcium-binding proteins [4]. They diversified in function through major evolutionary transitions, including the formation of a calcified skeleton [2], the emergence of teeth [5], and the introduction of lactation in mammals [6]. Many members of this gene family are relevant to oral biology as they encode not only proteins that form tooth enamel but also others that rank among the most abundant proteins in human saliva and participate in the protection and remineralization of tooth enamel [7].

With regard to their amino acid composition, SCPP genes can be divided into two subfamilies that form two distinct gene clusters on chromosome 4 encoding D/E/S-rich and P/Q-rich proteins, respectively [3]. In humans, these two clusters are separated by 17 megabases, with the overall genomic synteny remaining conserved across mammals [8]. The D/E/S-rich SCPP genes code for five acidic proteins (DSPP, DMP1, IBSP, MEPE, and SPP1), which have functions in bone and dentin mineralization [9, 10]. The P/Q-rich locus comprises 16 previously described (not including the distal P/Q-rich *SCPPPQ1* gene), syntenically clustered, genes and one distal enamel-related gene, *AMEL*, located on the sex chromosomes. These P/Q-rich SCPP genes possess functions in the mineralization of the enamel matrix and calcium-binding in milk [5, 6]. Notably, 5 out of the 17 P/Q-rich proteins encoded in this locus are highly expressed in salivary glands and belong to the most abundant proteins secreted in human saliva [7] **(Table 1).**

**Table 1.** SCPP gene names, reported functions, and tissue expression trends. “Broad” expression is defined as previously found to be expressed across two or more tissues. Genes are shown on the left as boxes colored by their expression categories: purple, enamel; yellow, salivary glands (saliva); blue, mammary glands (milk); red, broad expression; orange, acidic SCPP genes, bone and/or dentin. This table represents results from previous work (legacy) from studies cited throughout the manuscript and provides a starting framework for the downstream analysis and discussion. The functions listed in this table are derived from Human Protein Atlas and GeneCards.

| Gene | Function | Expression |
| --- | --- | --- |
| **CSN1S1**: Casein Alpha S1 | calcium binding, nutrition source | lactating breast (milk) |
| **CSN2**: Casein Beta | calcium binding, nutrition source | lactating breast (milk) |
| **STATH**: Statherin | calcium binding, antimicrobial | salivary glands (saliva) |
| **HTN3**: Histatin 3 | calcium binding, antimicrobial | salivary glands (saliva) |
| **HTN1**: Histatin 1 | calcium binding, antimicrobial | salivary glands (saliva) |
| **CSN1S2**: Casein Alpha S1 | pseudogene | unknown |
| **PRR27**: Proline-Rich Protein 27 | unknown | broad |
| **ODAM**: Odontogenic, Ameloblast Associated | enamel mineralization | broad |
| **FDCSP**: Follicular Dendritic Cell Secreted Protein | immune regulation | broad (salivary glands) |
| **CSN3**: Casein Kappa | calcium binding, nutrition source | lactating breast (milk) |
| **CABS1**: Calcium Binding Protein, Spermatid Associated 1 | calcium binding, sperm flagella | broad |
| **SMR3A**: Submaxillary Gland Androgen Regulated Protein 3A | regulation of sensory perception of pain | salivary glands (saliva) |
| **SMR3B**: Submaxillary Gland Androgen Regulated Protein 3B | regulation of sensory perception of pain | salivary glands (saliva) |
| **PROL1**: Basic Proline-Rich Lacrimal Protein | regulation of sensory perception of pain | lacrimal and salivary glands (saliva) |
| **MUC7**: Mucin 7 | microbial binding, regulation of physical properties of saliva | salivary glands (saliva) |
| **AMTN**: Amelotin | enamel mineralization | ameloblasts |
| **AMBN**: Ameloblastin | enamel mineralization | ameloblasts |
| **ENAM**: Enamelin | enamel mineralization | ameloblasts |
| **AMEL**: Amelogenin (x-linked) | enamel mineralization | ameloblasts |
| **SCPPPQ1**:SCPP Proline-Glutamine Rich 1 | antimicrobial | ameloblasts |
| **SPARCL1**: SPARC Like 1 | calcium/metal-binding | broad |
| **DSPP**: Dentin Sialophosphoprotein | calcium-binding, dentinogenesis | odontoblasts |
| **DMP1**: Dentin Matrix Acidic Phosphoprotein 1 | dentinogenesis | osteoblasts and odontoblasts |
| **IBSP**: Integrin Binding Sialoprotein | bone mineralization | osteoblasts |
| **MEPE**: Matrix Extracellular Phosphoglycoprotein | bone mineralization, phosphate regulation | osteoblasts |
| **SPP1**: Secreted Phosphoprotein 1 | bone mineralization, signaling | broad (osteoblasts) |

In our previous work, we hypothesized that genes encoding proteins secreted by the salivary glands are more often subject to selective pressure due to arising differences in species-specific diets and habitats [11–13]. The P/Q-rich SCPP gene locus offers an excellent model to investigate this hypothesis. The locus harbors genes that code for some of the most abundant salivary proteins, yet these are clustered with evolutionarily related genes that code for proteins expressed in different tissues and exert fundamentally different functions. Since previous studies already focused on the involvement of SCPP genes in tooth mineralization [14] and calcium binding in milk [6], here we investigated the less-understood evolution of the P/Q-rich SCPP subfamily of genes, focusing on their function and evolution in saliva. Specifically, we set out to test the hypothesis that, within the limits of the P/Q-rich SCPP locus, saliva-related genes in mammals evolved faster than the neighboring enamel-related or milk-related genes. To achieve this goal, we conducted a bioinformatic examination across a wide range of vertebrate species, spanning from fishes to humans, to better resolve the evolution of the P/Q-rich SCPP gene locus. Given that the SCPP genes code for some of the most abundant salivary proteins in primates, including humans [7, 15], the insights gained from this study will ultimately enhance our understanding of how mammalian saliva co-evolved with species-specific habitat-driven diet preferences and environmental pathogen challenges.

## Results

### SCPP genes with novel functions are derived from conserved enamel-related genes

Gene gains and losses recurring in a given locus are rare across the genome [16]. The SCPP gene locus was pointed out in a review of genome-wide evolutionary trends as one of several examples where genes undergo a rapid turnover [17]. To find out how different functional categories of SCPP genes are affected by this rapid turnover, we analyzed the genetic variation in the SCPP locus in 61 vertebrate species comprising mammals (n=48), birds (n=4), reptiles (n=4), amphibians (n=1), and fishes (n=5) (**Figure 1**). Our study builds upon prior studies, which linked the variation in this locus to calcification of skeleton and teeth, as well as lactation [5, 6, 18]. The availability of a greater number of well-annotated vertebrate genomes allowed us to probe for the presence or absence of human SCPP orthologs and homologs in other vertebrates. We used a three-pronged approach: (i) searching for sequence similarity, (ii) comparing gene order, and (iii) analyzing gene models available on RefSeq and Ensembl. This analysis allowed us to revisit the conclusions previously made by Kawasaki [19] and tie them together with our new analyses to provide an updated and refined history of structural changes in the SCPP cluster of genes.

**Figure 1:**
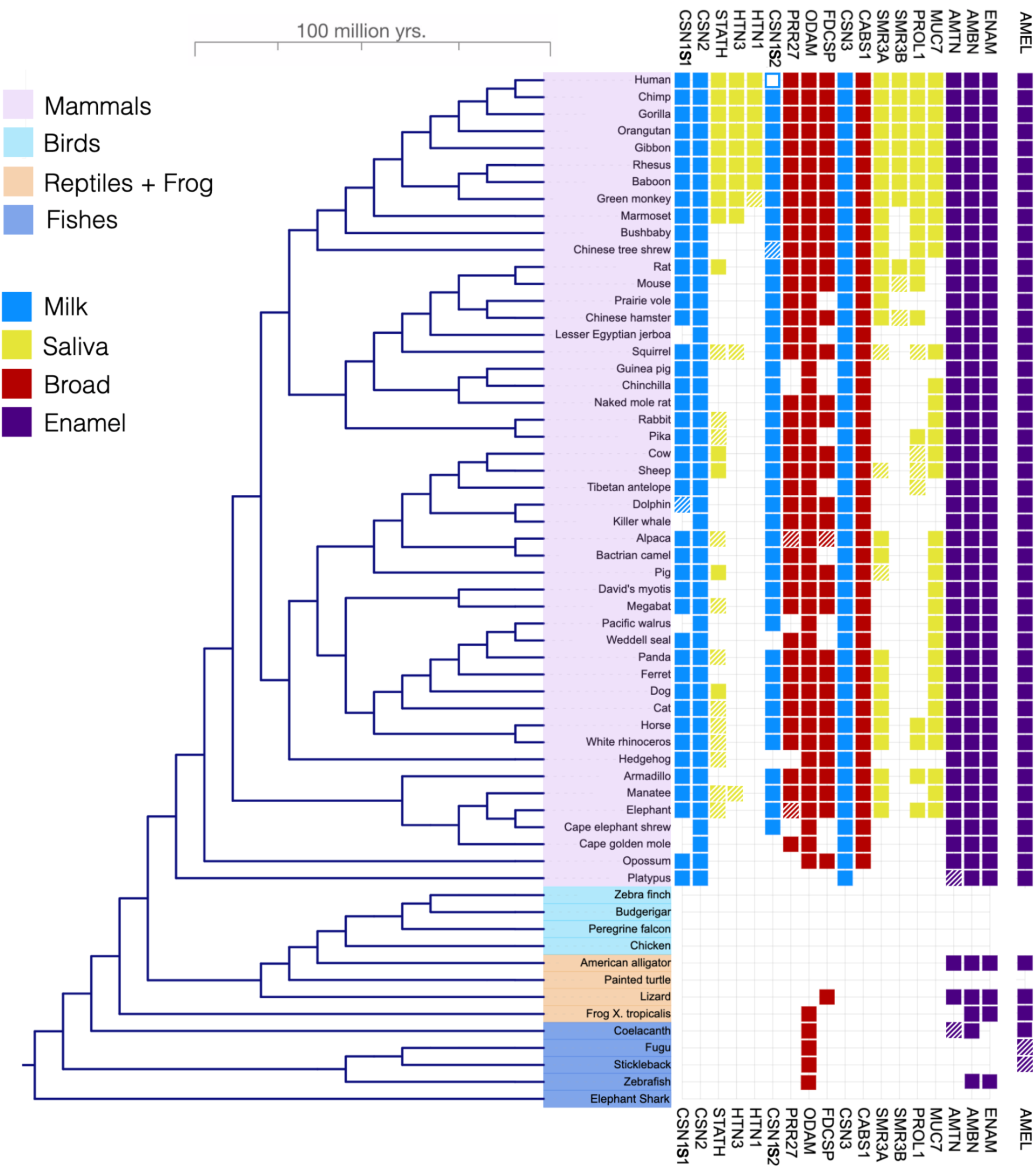
P/Q-rich SCPP genes in the vertebrate phylogeny. The presence and absence of orthologues to the P/Q-rich SCPP genes found in humans are indicated within a phylogeny of 61 vertebrate species comprising 48 mammals, 4 birds, 3 reptiles, 1 amphibian, and 5 fishes. Genes are shown as boxes with colors indicating expression categories: purple, enamel; yellow, salivary glands (saliva); blue, mammary glands (milk); red, broad expression (broad). Solid-colored boxes represent the presence of SCPP genes consistent between RefSeq and Ensembl databases. Hatched boxes indicate genes found in only one of the two databases.

Previous studies provided evidence that the tooth enamel-related SCPP genes *ENAM, AMBN,* and *AMTN* likely originated in fish ancestors [19, 20]. Our phylogenetic analysis corroborates earlier findings [21, 22], including the debated convergent evolution of enamel-related genes [23, 24] (**Figure 2**). We found that the presence of enamel-related SCPP genes correlated with the existence of teeth (**Figure 1)**. We further showed that all mammals included in our analysis harbor the enamel-related SCPP genes. These enamel-related genes, which originated when teeth evolved in fishes, were then conserved in reptiles and mammals and became the ancestral basis for the expansion of the wider SCPP gene family in mammals.

**Figure 2:**
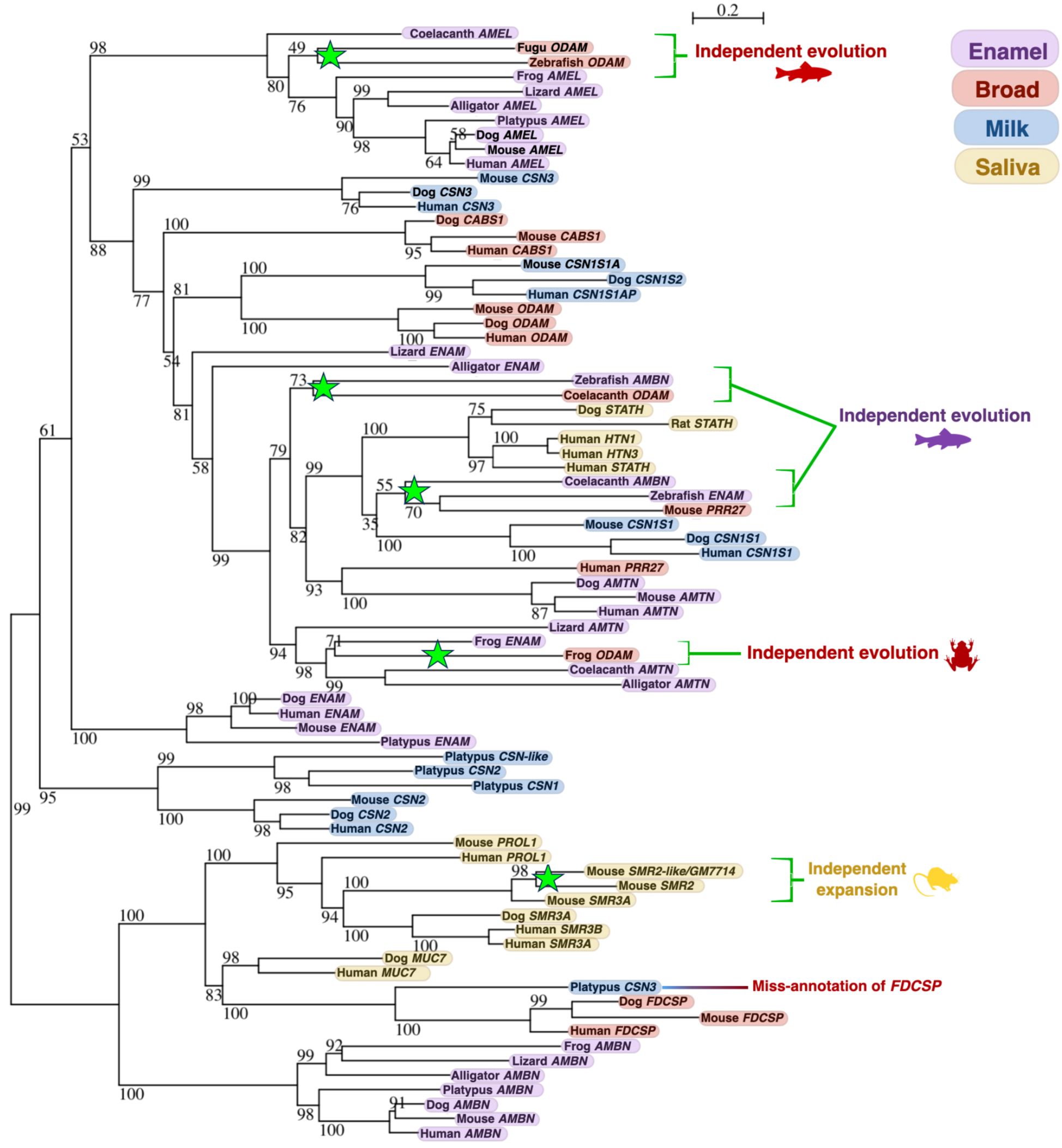
Phylogenetic tree of SCPP genes. Maximum likelihood tree generated using coding nucleotide sequences from major clades [mammals (human, mouse, dog, platypus), reptiles (lizard, alligator), an amphibian (frog), and fishes (coelacanth, fugu, zebrafish)]. Bootstrap support is shown at the nodes. Genes are shown at the leafs of the tree with colors indicating expression categories: purple, enamel; yellow, salivary glands (saliva); blue, mammary glands (milk); red, broad expression (broad). Green stars indicate the likely independent duplication events.

### Ancestral P/Q-rich SCPP genes in mammals gained broad-ranging functions, priming the locus for evolutionary innovation

The evolutionary mechanisms through which the ancestral enamel-related genes duplicated to give rise to other SCPP genes critical for salivary and mammary function are not well understood. We found that *ODAM, CABS1, PRR27*, and *FDCSP*, which all encode proteins rich in the amino acids proline and glutamine (P/Q-rich), are conserved in all mammals and, in some cases, reptiles (**Figure 1**). However, in contrast to the enamel-related SCPP genes, they show broad expression patterns (two or more tissues) (**Supplementary Figure 1**).

*ODAM* was proposed to be the predecessor to the other P/Q-rich SCPP genes and was thought to have evolved from *ENAM* in the bony fish ancestor [25]. Our data is concordant with *AMBN* being a possible ancestor of P/Q-rich SCPP genes. However, our analysis suggests that the origin of *ODAM* is more complicated than that. The evidence for this hypothesis is as follows: First, we found two genes, one in bony fish and one in mammals, both annotated as *ODAM*, but located in syntenically distinct genomic regions as described previously [26]. Second, a nucleotide-level phylogeny of *ODAM* shows two clusters, supporting that two independent duplication events took place (**Figure 2**). Third, the fact that these evolutionarily distinct genes that differ at the nucleotide level were both annotated as *ODAM* may stem from the fact that they are similar at the amino acid sequence level (**Supplementary Figures 2 & 3**). Thus, our findings suggest that genes annotated as *ODAM* may have evolved through adaptive convergence.

The *FDCSP* gene was proposed to have evolved from *ODAM* [5]. However, we found that *FDCSP* may have evolved earlier through a duplication of *AMBN.* Based on sequence similarity and synteny, we propose that a gene annotated as *CSN3* in the platypus genome may be miss-annotated and is, in fact, *FDCSP* (**Figure 2, Supplementary Figure 4**). If so, this observation places *FDCSP* as an ancestral gene from which other P/Q-rich SCPP genes in mammals have evolved (**Figure 3**). The *FDCSP* gene, itself expressed in a variety of different tissues (**Table 1**, [27]) may have given rise to PQ-rich genes with more specialized functions in saliva and milk, which is a typical example of subfunctionalization [28]. Interestingly, our analysis revealed two additional P/Q-rich genes in the locus (*PRR27* and *CABS1*) that had not been previously described as belonging to the SCPP gene family.

**Figure 3.**
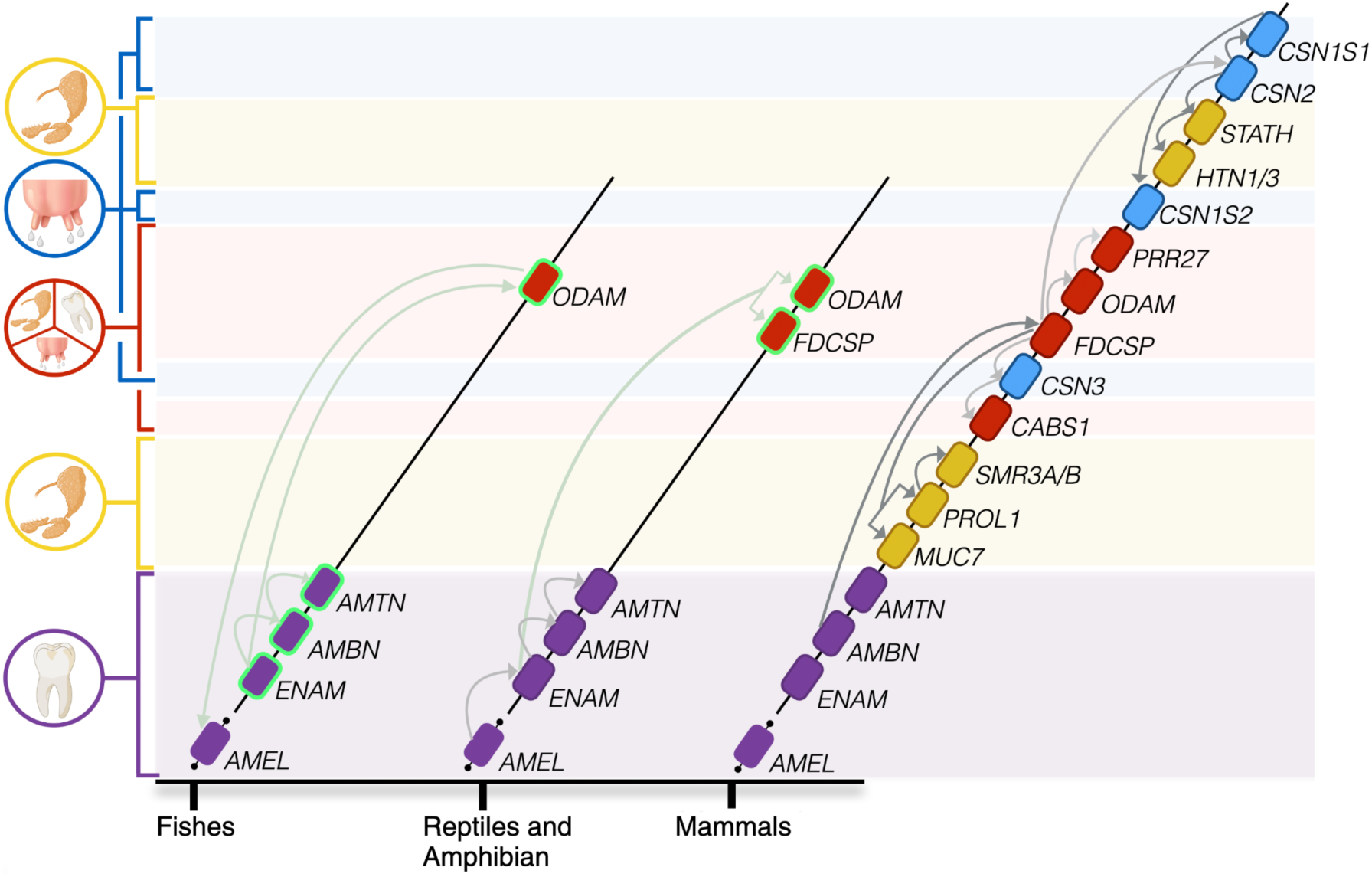
SCPP gene repertoire and evolutionary history in major vertebrate clades. The x-axis shows the three major vertebrate clades; fishes, reptiles/amphibians, and mammals. Birds were not included because no P/Q-rich SCPP genes were found. Genes are shown as boxes with colors indicating expression categories: purple, enamel; yellow, salivary glands (saliva); blue, mammary glands (milk); red, broad expression (broad). Genes that we proposed to have arisen independently, but were annotated with the same name, are outlined in green. Arrows show the proposed origins of genes within the SCPP gene family. Darker arrows indicate higher confidence in the origins, and lighter lines are cases where weaker evidence was found for the origin of duplication. The “/” symbol is used to group paralog genes that are highly similar in sequence, and have been further resolved in a more thorough analysis of the primate gene locus (see below and Supplementary Figure 6)

### Divergence of the milk-related casein family of genes through independent, recurrent, and subsequent duplication events

The milk-related caseins encoded in the SCPP locus are major components in mammalian milk, with an ability to bind calcium [29], thereby ensuring that this nutritive biofluid remains supersaturated with calcium for aiding skeleton formation in growing nurslings. In mammals, they form two casein types: calcium sensitive caseins (*CSN1S1, CSN1S2A*&*B, CSN2)* and calcium-in-sensitive caseins (*CSN3*) [6]. Their evolutionary history and diversity may be linked to the evolution of lactation in different mammalian species. Kawasaki and coworkers [6] proposed that the calcium-sensitive casein types evolved before the emergence of mammals through duplication of *SCPPPQ1* (another SCPP gene thought to have originated from *ODAM* [6]) that translocated into the P/Q-rich SCPP locus. Focusing on the evolution of caseins within the P/Q-rich SCPP locus, we showed close sequence similarity between the *FDCSP* gene and caseins, indicating that *FDCSP* is ancestral to caseins. Further supporting this notion, we found that what is annotated as *CSN3* in platypus should be annotated as *FDCSP*, based on the phylogenetic clustering of this gene (**Figure 2**). Specifically, a parsimonious reading of the phylogenetic clustering suggests that the ancestral caseins *CSN3* and *CSN2* may have evolved through independent tandem duplications of the *FDCSP* gene (**Figure 3**, **Supplementary Figure 4**). This aligns with previous work [6, 7] suggesting independent origins of different casein types, however, our results highlight that *FDCSP* may have given rise to both types of caseins. This is supported by the fact that *ODAM* had not yet evolved to give rise to *FDCSP* nor to *SCPPPQ1* in mammals.

Within the casein subfamily of genes, we found that beta-casein (*CSN2*) and kappa casein (*CSN3*) genes were conserved in all mammals investigated, which is concordant with other studies [30]. In contrast to that, we found that the alpha caseins (*CSN1S1* and *CSN1S2*) were less conserved and varied in different mammalian lineages. Notably, *CSN1S1* is absent in some marine mammals (**Figure 1**), and has gained a loss of function mutation in the catarrhini (Old World monkeys and great apes) ancestor, making it a pseudogene in multiple primates, including humans (**Supplementary Figure 5**). Combined, our study expands on previous work [6], demonstrating that different mammalian species have evolved diverse casein repertoires, perhaps shaping the various lactation strategies and nutritional content of the milk across mammals.

### Varied origins of saliva-related SCPP genes in primates

Orthologs of human saliva-related SCPP genes that are expressed in salivary glands and secreted into saliva [7] were found to be most dynamic in their presence or absence among nonhuman mammalian species (**Figure 1**). Even though previous studies investigated the presence and absence of some of these proteins in the saliva of different species [31], their evolutionary history remained unclear. Our results suggest three groups within human salivary SCPP genes that seem to have a distinct evolutionary history.

First, we investigated the evolution of *STATH* as it was argued that *STATH* duplicated from a casein gene [4] *CSN1S2,* and gave rise to the primate-specific histatin (*HTN*) genes [32]. The analysis of gene duplication history is complicated by potential gene conversion or fusion of duplicated segments within the locus. Having said that, phylogenetic clustering suggests that *CSN1S1,* as opposed to *CSN1S2,* is the more likely progenitor of *STATH* (**Figures 2** and **3**) and that *STATH* has evolved in the primate lineage after divergence from the tree shrews (**Supplementary Figure 6**). It is of note that we found *STATH*-like sequences within the SCPP locus in non-primate species (**Figure 1**). This can be explained by multiple losses of this gene across the phylogeny. However, we think a more parsimonious explanation would be that convergent evolution of the signal peptide portion of this small protein in non-primate mammals from other SCPP genes may account for what we observed (**Supplementary Figure 6)**. For example, in cows, there are two histatin-like genes annotated as *STATH* and *HSTN* (in cows called histatherin) with very similar signal peptide sections, but observably different amino acid sequences from each other and the canonical primate *STATH* genes. Future work based on more complete reference genomes may resolve this issue. Functionally, we propose that *STATH* retained the original casein function of preventing calcium precipitation, but executes this function in saliva now instead of in milk. Furthermore, we resolved that *HTN3* was duplicated first from *STATH* **(Figures 2 and 3)**, followed by the duplication of *HTN1* from *HTN3* in the ancestor of catarrhini (**Supplementary Figure 6**). The histatins did not retain the calcium-binding function of caseins but were proposed to have evolved zinc- and copper-binding motifs instead [33].

*SMR3A* and *SMR3B*, along with *PROL1* (also named *OPRPN*), comprise a group of salivary SCPP genes that code for proteins rich in proline and have a distinct phylogenetic history. We previously showed that these proteins, due to their richness in proline, have served as precursors for the *de novo* evolution of mucin function at various stages of the mammalian phylogeny [13]. Our current analysis confirmed that based on nucleotide sequence homology, *SMR3A* and *SMR3B* likely originated as duplicates of *PROL1* [34]. The origin of *PROL1* itself goes back to the ancestor of placental mammals (**Figure 1**), but a possible precursor to *PROL1* with its short and repetitive sequence remains elusive.

*MUC7*, a short soluble mucin that neighbors *PROL1* in the human genome, is expressed abundantly in salivary glands but has no clear evolutionary connection to other salivary SCPP genes. Our analysis here confirmed our previous assertion [11] that *MUC7* evolved in the placental mammal ancestor. It was previously suggested that *MUC7* might be evolutionarily related to *STATH* and *HTNs* because it harbors a histatin-like domain in its N-terminus [35]. However, we found that *MUC7* phylogenetically clusters with genes encoding proteins rich in proline (**Figure 2**). Collectively, *MUC7* evolution has been complex and likely involved gene duplications and fusions [11, 36], as well as the recurrent acquisition of exonic repeats encoding proline, threonine, and serine rich O-glycosylated mucin domains [13].

### The majority of SCPP genes expressed in human salivary glands are primate-specific

Based on the absence of multiple saliva-related SCPP genes in non-primate vertebrates (**Figure 1**), we asked whether the salivary gland tissue-specific expression of SCPP genes differs between primates and other mammals. Since expression data of all three major salivary gland tissues in other mammals are available only for mice [37] and humans, we document the expression of SCPP genes in human tissues (**Supplementary Figure 1**) and compared SCPP gene expression between human and mouse salivary glands (**Figure 4**).

**Figure 4:**
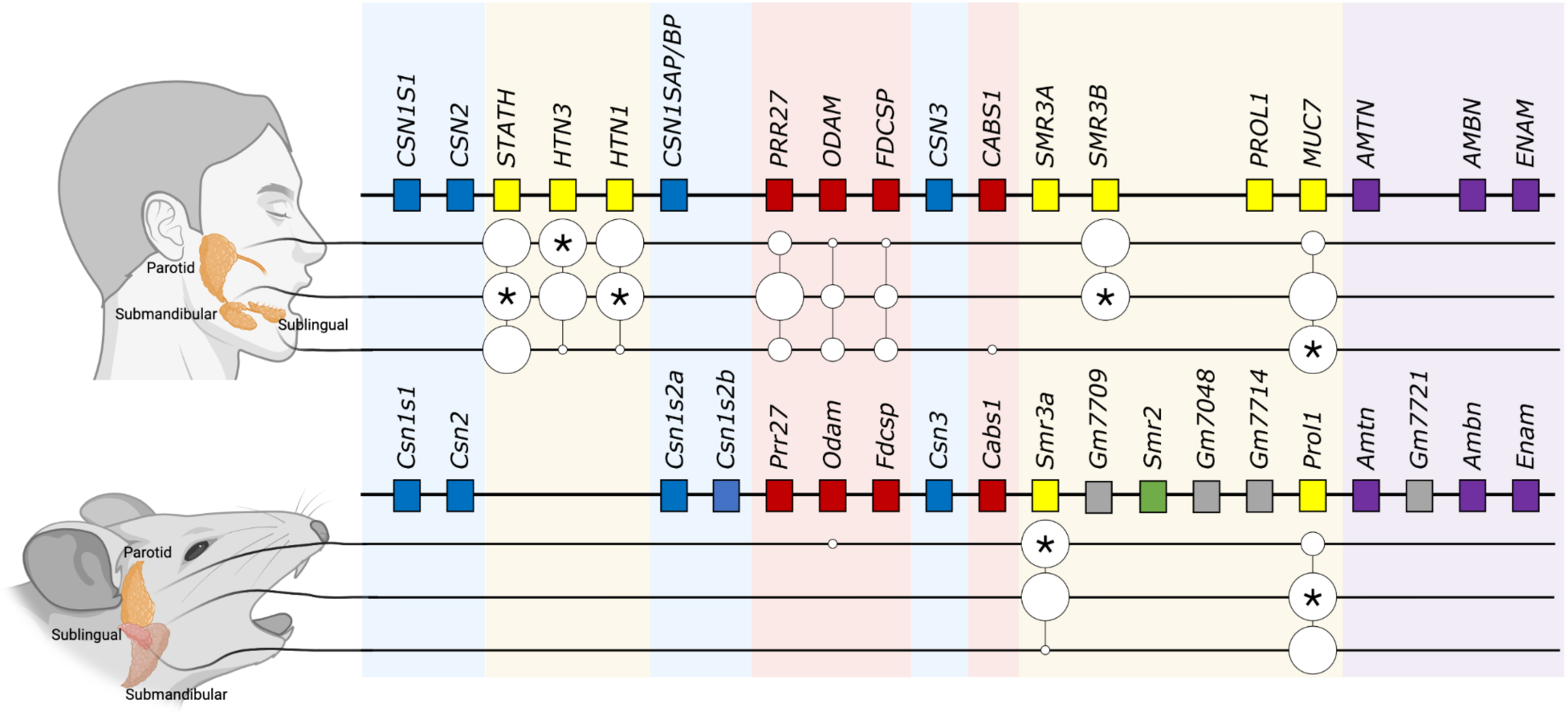
SCPP gene turnover and shifts in tissue expression. Comparison of SCPP gene expression in the three major salivary gland types between human and mouse. SCPP genes are depicted as boxes and colored based on expression categories: purple, enamel; yellow, salivary glands (saliva); blue, mammary glands (milk); red, broad expression (broad). The mouse-specific Smr2 gene is indicated in green, and four gene duplicates, that are annotated in the mouse reference genome as pseudogenes, are indicated in grey. The upper part of the panel shows the three major human salivary glands (parotid, submandibular, and sublingual). The lower part of the panel shows the mouse salivary gland counterparts. The size of the bubbles indicates the expression level categorized as low, medium, or high. For genes with high levels of expression, the glands having the highest expression are indicated by stars. Genes are arranged in how they appear in the synteny of human and mouse reference genomes. Transcriptional data for human and mouse salivary gland tissues were obtained from previously published reports [7, 37].

We found that, except for *SMR3A*, none of the SCPP genes that are expressed in human salivary glands are expressed in the mouse salivary glands (**Figure 4**). This is an unexpected finding, given that the most highly expressed genes are conserved and retain expression trends across species [38, 39]. The expression differences we found can be explained by the absence of *HTN1*, *HTN3*, *STATH*, and *MUC7* in the mouse genome and by the only minimal expression of existing orthologs of human saliva-related genes (*PRR27*, *FDCSP*, and *ODAM*). Besides *SMR3A*, the only abundantly expressed SCPP gene in the mouse salivary glands is the lineage-specific gene, *MUC10*, which has evolved from *PROL1* by acquiring O-glycosylated mucin PTS repeats [13]. In sum, our results show that a combination of gene turnover events (loss and emergence of genes) and putative changes in regulatory sequences led to remarkable differences in the salivary expression of mouse and human SCPP genes.

### Saliva-related SCPP genes in primates evolved faster than milk- and enamel-related ones

We showed that the evolutionary origins of saliva-related SCPP genes in primates are more recent than those of enamel-related and milk-related genes (**Figure 1**). To expand on this observation, we next investigated nucleotide level variation in P/Q-rich SCPP genes. By applying conservation scores to our species phylogeny, we find that enamel-related and milk-related genes, along with the broadly expressed genes, are significantly more conserved than the saliva-related SCPP genes (p<2.2 x 10^-16^, **Figure 5, panels a** & **b**). Our observation is consistent with a scenario where enamel-related SCPP genes have evolved under purifying selection, as previously suggested [40], while saliva-related genes have experienced bouts of lineage-specific selection.

**Figure 5:**
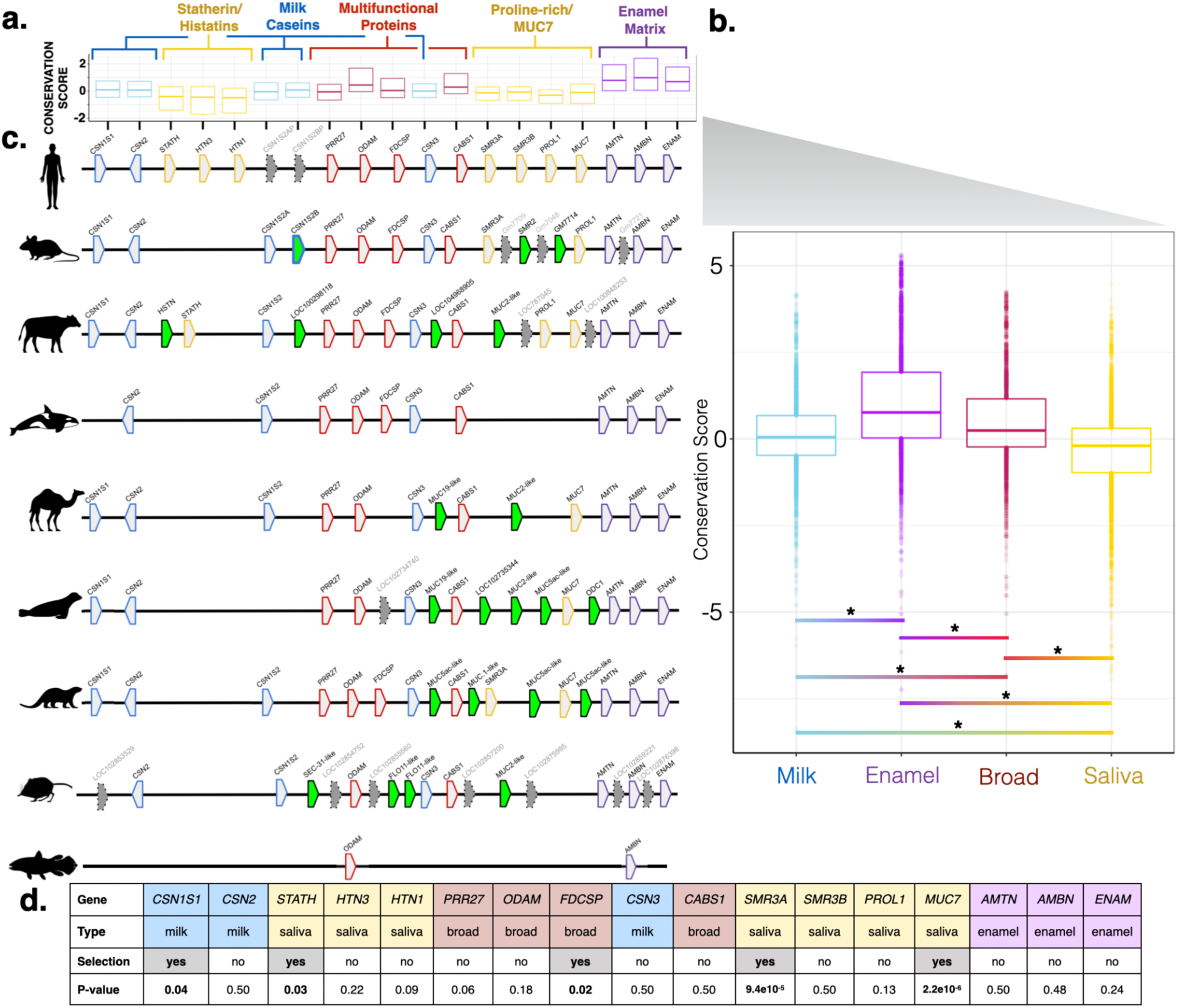
Gene conservation and turnover within the SCPP locus. **(a)** Box plots represent conservation scores calculated for different SCPP genes across 100 vertebrate species (phyloP100way) for each SCPP gene on a per-base-pair basis. The boxes represent the average gene conservation score (+/- 2 standard deviations). Colors denote the functional category: purple, enamel; yellow, salivary glands (saliva); blue, mammary glands (milk); red, broad expression (broad). **(b)** Box plot representing conservation scores calculated per base pair for the P/Q-rich SCPP genes but merged by functional category. Stars (*) indicate significance scores p<2.2×10^-16^, and bar colors show comparisons for respective categories. **(c)** The SCPP loci for nine selected species are shown to illustrate the turnover of genes within the locus. Genes are labeled according to their functional categories (same as in panel a). Lineage-specific genes are colored in green. Pseudogenes are colored in grey with dotted outlines. **(d)** Table summarizing the results of the selection analysis using BUSTED. Colors denote the category of function for each gene.

Given the diversity of diets consumed by different mammalian species, we hypothesized that the saliva-expressed SCPP genes evolved faster than their milk- and enamel-related counterparts in the P/Q-rich SCPP gene locus. Support for this hypothesis comes from the observation that almost half of the salivary SCPP genes in primates do not have orthologs in non-primate genomes, whereas most other genes in this locus have orthologs among primates and non-primate mammals (**Figure 1**). To visualize the rate of gene turnover among mammals, we show the SCPP genes in nine species for which high-quality genome assemblies are available and that exemplify major mammalian phylogenetic groups with unique habitats, ecological contexts, and different diets (**Figure 5c**). We found multiple lineage-specific genes in these mammals. Many of the lineage-specific genes were shown in our previous work to be so-called “orphan” mucin genes, which recurrently evolve in the P/Q-rich SCPP gene locus from precursor genes that code for proteins rich in proline and have a propensity to be expressed in saliva [13]. Overall, this observation opens the possibility that saliva-related genes in the P/Q-rich SCPP locus are proportionally more often affected by gene turnover events.

Based on the outcome of the gene conservation analysis, we hypothesized that lineage-specific directional selective forces affect the saliva-expressed SCPP genes more than the milk- and enamel-related genes in the locus. To test this hypothesis, we searched for lineage-specific selection acting on human SCPP genes and their orthologs in nonhuman primates. We focused our analysis on primates because the majority of saliva-related human SCPP genes do not have orthologs in non-primate mammals (**Figure 1**). Using BUSTED, a likelihood ratio test for diversifying selection affecting select branches in a given phylogenetic tree [41], we found evidence of lineage-specific selection for 3 out of 7 saliva-related P/Q-rich SCPP genes, but only for 2 out of 10 other P/Q-rich SCPP genes (**Figure 5d**). Additional evidence of selection acting on saliva-related SCPP genes was derived from dN/dS (nonsynonymous versus synonymous substitution) pairwise comparisons (**Supplementary Figure 7**). Our results support the hypothesis that primate-specific adaptive, rather than neutral, forces led to the accelerated evolution of saliva-related SCPP genes in primates. Collectively, our data identify the P/Q-rich SCPP locus, and, in particular, the saliva-related genes, as a hotbed of evolutionary adaptation, where structural, nucleotide, and regulatory differences shape the functional diversity of mammalian saliva.

## Discussion

### Expansions of different functional gene groups within the SCPP locus occurred during major evolutionary transitions

The expansion of particular SCPP genes appeared alongside major evolutionary transitions. The early origin of *SPARCL1* and the subsequent evolution of mineralized tissues is consistent with the hypothesis that the origin and diversification of vertebrate mineralized tissues are correlated with the origin and episodic duplications of SCPP genes [42]. The first expansion of gene diversity in the SCPP cluster took place along with the evolution of tooth enamel and created the genes *ENAM, AMBN,* and *AMTN*. The second expansion in the SCPP gene family occurred along with the evolution of lactation in mammals and affected the milk-related casein subfamily of genes. Here, we propose that a third, most recent expansion in the SCPP gene family affected the saliva-related SCPP genes in primates, including humans.

Generally, sequence conservation is the hallmark of biological function. However, a small number of loci with important functions may adaptively change in certain species due to lineage-specific adaptations. This process creates a paradox where functionally important loci will lose sequence conservation across species, creating what Ponting dubbed as “evolutionary twilight zones” [17]. In fact, he highlighted the SCPP locus as one such locus, making it a natural model to address fundamental questions regarding accelerated lineage-specific evolution. Focusing on the P/Q-rich SCPP gene locus, where enamel- and milk-related genes are lined up in the vicinity of saliva-related genes, we found that the saliva-specific SCPP genes underwent accelerated evolution in the primate lineage, both through gene duplications and amino-acid substitutions. The majority (5 out of 7) of saliva-related SCPP genes either emerged in primates or showed signatures of positive selection in the primate lineage. One of the outstanding questions, especially in light of a recent look into the regulatory sequences in this locus in mice [43], is how the regulatory architecture of this locus has evolved to maintain expression that is specific to salivary glands.

Since the expansion and diversification of saliva-related SCPP genes took place along with the diversification of the primate lineage, occupancy of different dietary niches, and exposure to different pathogenic challenges, we propose that this third expansion of genes within the SCPP locus could have been driven by primate species-specific selective pressures. Indeed, dietary change has been discussed as a major adaptive pressure in primates [44–47]. Despite the scarcity of clear functional data for saliva-related SCPP proteins, several lines of evidence support the hypothesis that saliva-related SCPP genes have evolved to accommodate the diverse dietary needs of primates, shaping taste perception [48, 49] and physical properties of saliva [50]. In parallel, numerous SCPP genes have been identified as key players in binding and interacting with bacteria and other pathogens in saliva [33, 51–58]. Within this context, it is conceivable that an ongoing evolutionary competition between pathogens and the saliva-related SCPP proteins partially underlies the observed SCPP genetic variation among primates. What we demonstrated here in primates suggests that evolutionary adaptations in saliva-related SCPP genes, particularly through gene losses, recurrent duplications, and alterations in regulatory sequences, may be critical also in other branches of the mammalian phylogeny for rapidly adapting to environmental pressures, including dietary diversification and pathogenic challenges.

## Conclusion

Our study explores the gene evolution in the P/Q-rich SCPP locus and helps understand the formation of diverse gene families. Our work clarifies the gene turnover and the sequence change in the SCPP locus within the context of three major evolutionary transitions corresponding to the emergence of teeth, the development of milk in mammals, and dietary diversification in primates. The accelerated evolution of saliva-related genes explains the observation that salivary proteomes vary significantly among primates, likely shaped by the environments and dietary habits of different species. What was shown here for primates, suggests that species within other phylogenetic orders, when adapting to diverse habitats, unique diets, or varying microbial encounters, may have undergone a similar evolutionary expansion of saliva-expressed genes. Further investigation of saliva proteomes and the corresponding transcriptomes of salivary glands among mammals in diverse ecological niches with distinct dietary specializations is needed to test this hypothesis.

## Methods

### Gene Tree and BLAST parameters

Species radiation and divergence parameters were downloaded from the UCSC Genome Browser: http://hgdownload.cse.ucsc.edu/goldenpath/hg19/multiz100way/hg19.100way.nh. Tree visualization was done using the program iTOL [59]. NCBI BLAST was used to determine the presence or absence of genes in reference genomes of species. Human nucleotide and mRNA sequences were downloaded from Ensembl. Human protein sequences were downloaded from UniProt [60]. We also utilized SCPP gene sequences from the mouse genome with the same gene names as in humans to interrogate these loci in other species. FASTA protein sequences were first queried in each individual species using blastp (non-redundant protein sequences). We then searched nucleotide sequences in parallel using tblastn option on NCBI BLAST. Scoring parameters for the blastp [61] algorithm were chosen as follows:

matrix: BLOSUM62,

gap costs: existence 11 extension 1,

compositional adjustments: composition score matrix adjustment [62].

BLAST hits were assessed based on max score, total score, query cover (>30%), E-value (<0.01) and identity percentage (>20%). Top hits were manually inspected using respective gene models on the species’ reference genomes as well as reciprocal BLAST confirmation. The amino acid sequences of SCPP genes were then downloaded and mapped back to the human reference genome (Hg19) using the BLAT tool on UCSC genome browser to confirm relative homology. All SCPP gene inconsistencies from human reference were confirmed by examining individual gene models. To clarify any BLAST hits that are outside of our target significance level, we manually checked gene order across species using gene models provided by NCBI available through UCSC Genome Browser. The two highly conserved genes sulfotransferase family 1E (*SULT1E1*), and immunoglobulin J chain; (*JCHAIN*), flank the SCPP locus and thus were used to demarcate the SCPP chromosomal location and gene synteny. As a supplement, UCSC Genome Browser [63] gene predictions were used to confirm and contrast gene presence or absence. One major challenge in our analysis is that most salivary SCPP genes are short and encode proteins rich in proline, thus harboring repetitive codons. Thus, the power to identify sequence homology across species and between paralogs is lower than for larger genes with less repetitive sequence content.

Our approach identified examples of potentially erroneous gene annotations and other potential complications. Since *FDCSP* and the casein genes have some sequence homology resulting from duplication and *CSN3* is syntenically adjacent to *FDCSP*, it results in a misannotation. As such, the *FDCSP* gene is miss-annotated as *CSN3* in the platypus. In some cases, genes were annotated as a “LOC”, typical of inconclusive annotation. Further, many species exhibit species-specific independent duplications of SCPP genes, such as *CSN1S2* in rodents, and *SMR*-like gene expansion in mice. It is important to note that there may be other pseudogenization events of certain SCPP genes in a species-specific manner that were not detected in our study.

### Phylogenetic Trees

Amino acid and coding nucleotide sequences were downloaded from the RefSeq database for major clades [mammals (human, mouse, dog, platypus), reptiles/amphibian (lizard, alligator, frog), and fishes (coelacanth, fugu, zebrafish)]. Sequences were aligned using Clustal Omega in Seaview. Maximum likelihood trees were constructed using IQtree 2 [64] with 1000 bootstrap replicates. We did not include gaps in our calculations. The resulting data were used to generate **Figure 2** and **Supplementary Figure 3** Simplified trees with enamel-related genes were used in **Supplementary Figure 2**. We categorized events as independent when the nucleotide-based phylogeny showed distinct phylogenetic clustering. We found at least one example of possible convergent evolution in ODAM, where nucleotide-based phylogeny shows independent evolution in multiple branches, while amino-acid sequence clusters in a single branch. Sequence alignments can be found in **Additional File 1**.

### Duplication History

The duplication history shown in **Figure 3** was constructed using several criteria. Sequences were downloaded as mentioned above. Sequences of different SCPP genes in different species (both nucleotide and protein, respectively) were aligned using Seaview, and phylogenetic trees were constructed as described above (**Figure 2, Supplementary Figure 2 & 3**). Combining information for the clustering of genes with high bootstrap support values allowed us to ascertain a potential sequential duplication history, albeit confounded by independent gene gain and loss events. It is important to note that it is difficult to distinguish between the parent and an offspring gene. This analysis is further complicated by the divergence times of analyzed species and the genome qualities of many of the species assessed. Thus, we considered replication of previously described duplication events as confident and described in the text as such.

### Selection and Conservation Score Analysis

Positive selection analysis was performed using BUSTED [41]. Positive selection, which beneficial mutations increase in allele frequency in a population. In this context, BUSTED analyzes a DNA sequence alignment across multiple species and identifies positive selection by detecting sites within the sequence that have changed more than expected by neutral expectations of positive selection. BUSTED compares the rates of synonymous (silent) and non-synonymous (amino acid changing) substitutions at each site. Sites that have experienced positive selection will have a higher rate of non-synonymous substitutions than synonymous ones. BUSTED is a Bayesian-based method that incorporates prior knowledge about the distribution of selection pressures across different sites in the gene. The program uses a statistical model to estimate the proportion of sites under positive selection and the strength of selection at each site. Sequence alignments for primate species for each SCPP gene and their orthologs in nonhuman primates were downloaded from multiz100way [65] in the UCSC Genome Browser. We focused our analysis on primates because most human SCPP genes do not have orthologs in non-primate mammals. In addition, using MEGA X [66], we calculated global dN/dS (the ratio of substitution rates for nonsynonymous and synonymous SNPs) (**Supplementary Figure 7**) in a pairwise fashion, for the same primate species and gene alignments. dN and dS ratios were normalized by dividing the number of nonsynonymous SNPs by the human protein length, and the number of synonymous SNPs by human transcript length respectively. Verified protein lengths were extracted from the Uniprot database [60], while the longest protein-coding transcript lengths were obtained from the Ensembl database [67]. We used different databases because nucleotide alignments between species may be imperfect and introduce frameshifts. Site-by-site, pairwise, conservation scores were downloaded from the UCSC Genome Browser using the 100 vertebrates Basewise Conservation by PhyloP (phyloP100way) for each SCPP gene (**Figure 5**).

### Gene Expression Analysis

RNA sequencing data used to construct **Supplementary Figure 1** were taken from previous literature, salivary gland expression data from Saitou *et al.* [7], and GTEx [68] expression data from multiple tissues. Expression data for enamel-related genes was extracted from previous work [69]. The RNA sequencing data used to construct **Figure 4** was taken from [7] for human salivary glands, and from [37] for mouse salivary glands. To normalize between datasets, we ranked gene expression values by percentile in each tissue and based on that decided on below level of significance, low, medium, and high expression categories. The relative expression levels for mice, which is calculated as Fragments Per Kilobase of transcript per Million mapped reads (FPKM): high is >500, intermediate is <500 and >100, low is <100 and >25, and below level of significance is <25. The relative expression levels for humans, which are calculated in transcripts per million (TPM): high is >10,000, intermediate is <10,000 and >1,000, low is <1,000 and >100, not detectable is <100.

## Figures and Statistics

All data and figures were created using ggplot2 in R studio, Biorender, and the Keynote application. Statistical significance in Figure 4**, panels b** and **d** was calculated using the Wilcoxon rank sum test. In **Table 1**, the functions are derived from Genecards [70] and Human Protein Atlas [71].

## Acknowledgments

We thank Vincent Lynch for technical help, discussions, suggestions, and feedback on the manuscript. We thank Benjamin Stein for additional bioinformatic validation of the presence and absence of genes in species’ genomes.

## Funding

This study was funded by NSF grant no. 2049947 (to O.G.) and in part by National Institute of Dental and Craniofacial Research (NIDCR) grants R01 DE019807 and R21 DE025826 (to S.R.), and National Cancer Institute (NCI) grant U01 CA221244 (to S.R.).

## Author contributions

P.P., S.R., and O.G. conceived and designed the study and wrote the paper. P.P. performed all computational analyses. L.L. analyzed RNA-sequencing data for salivary glands in humans and mice and analyzed RNA-seq data in different organs and tissues in humans. All authors edited and approved the final version of the manuscript.

## Competing interests

The authors declare that they have no competing interests.

## Notes

### Competing Interest Statement

The authors have declared no competing interest.

