## Supplementary Figures for "Emergence of saliva protein genes in the secretory calcium-binding phosphoprotein (SCPP) locus and accelerated evolution in primates"

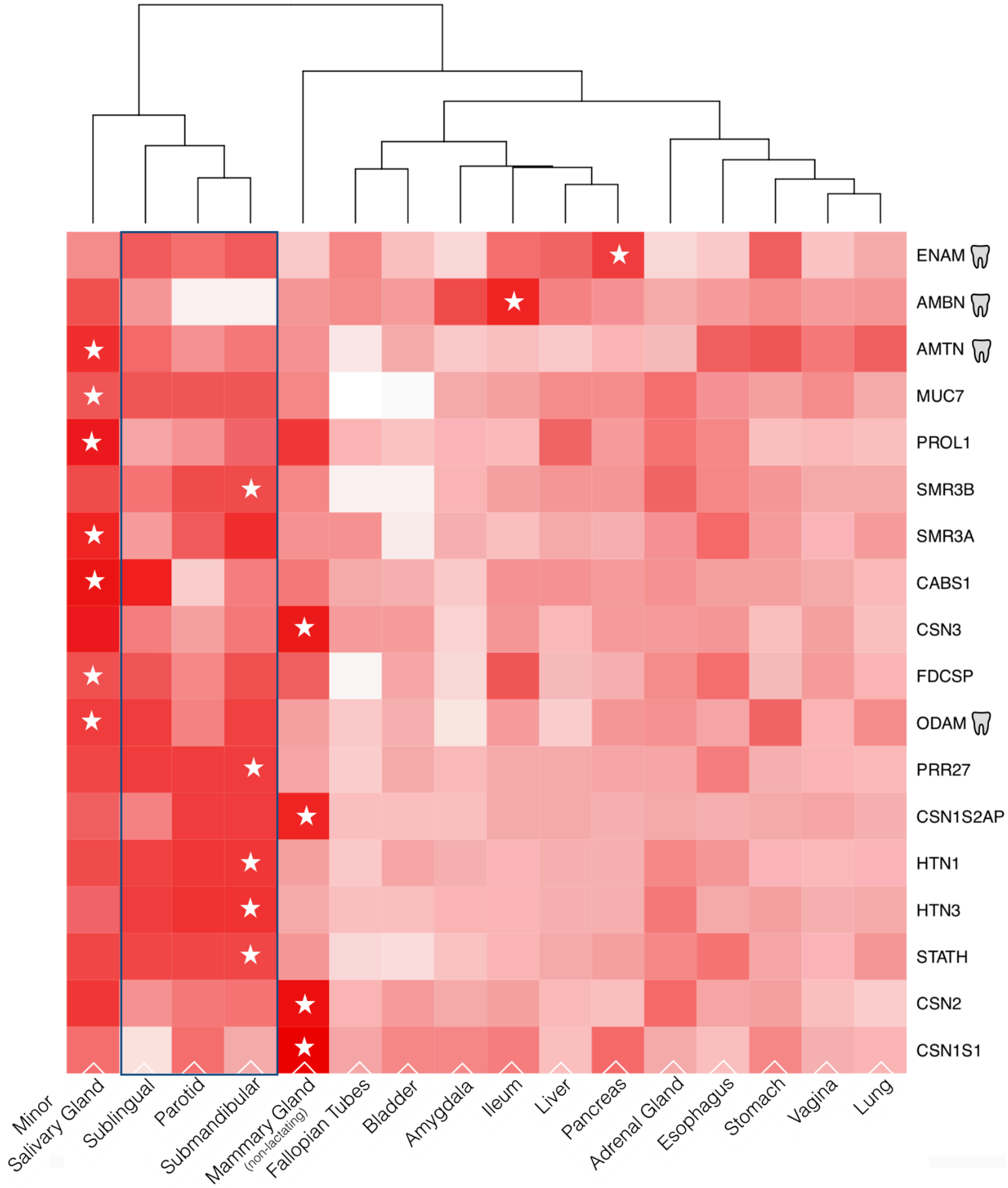


**Supplementary Figure 1: Expression of P/Q-rich SCPP genes across human tissues reveals they are mainly expressed in salivary and mammary glands.** The heat map shows relative gene expression within tissues from GTEx. Gene expression was ranked by percentile of expression compared to all genes found to be expressed in that tissue. More highly expressed genes are colored darker. The x-axis lists the tissues analyzed. Expression data of salivary gland tissues were obtained from our previous work [7] and indicated by dark blue outline. The y-axis lists the SCPP genes in the order that they appear in the genome (top is 3' end, bottom is 5' end). Tissues have been hierarchically clustered based on expression trends. Stars indicate the tissue where the gene was highest expressed. Tooth pictograms indicated genes where high expression was found in ameloblasts (enamel, not among the tissues of GTEx [69].

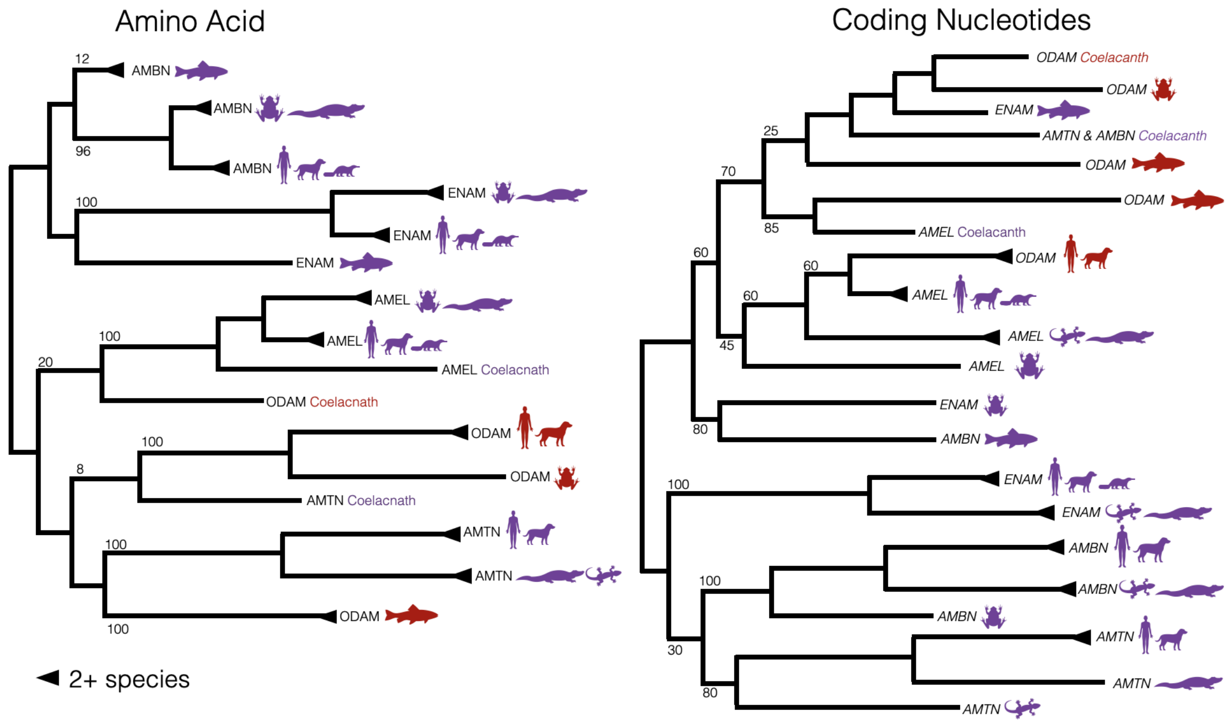


**Supplementary Figure 2: Maximum likelihood phylogenies show independent evolution and convergence of SCPP genes involved in tooth enamel mineralization and the corresponding proteins** . Phylogenetic trees were based on the amino acid (left) and nucleotide alignment (right) of *ODAM, AMTN, AMBN, ENAM,* and *AMEL*. Coding sequences and peptides were extracted for major clades (mammals [human, mouse, dog, platypus], reptiles [lizard, alligator], an amphibian (frog), and fishes (coelacanth, fugu, zebra fish)]. Species pictograms are shown and colored according to functional properties of enamel-related genes (purple) and broadly-expressed genes (red). Triangles indicate collapsed branches of two or more subclades. Bootstrap support values are indicated on branches.

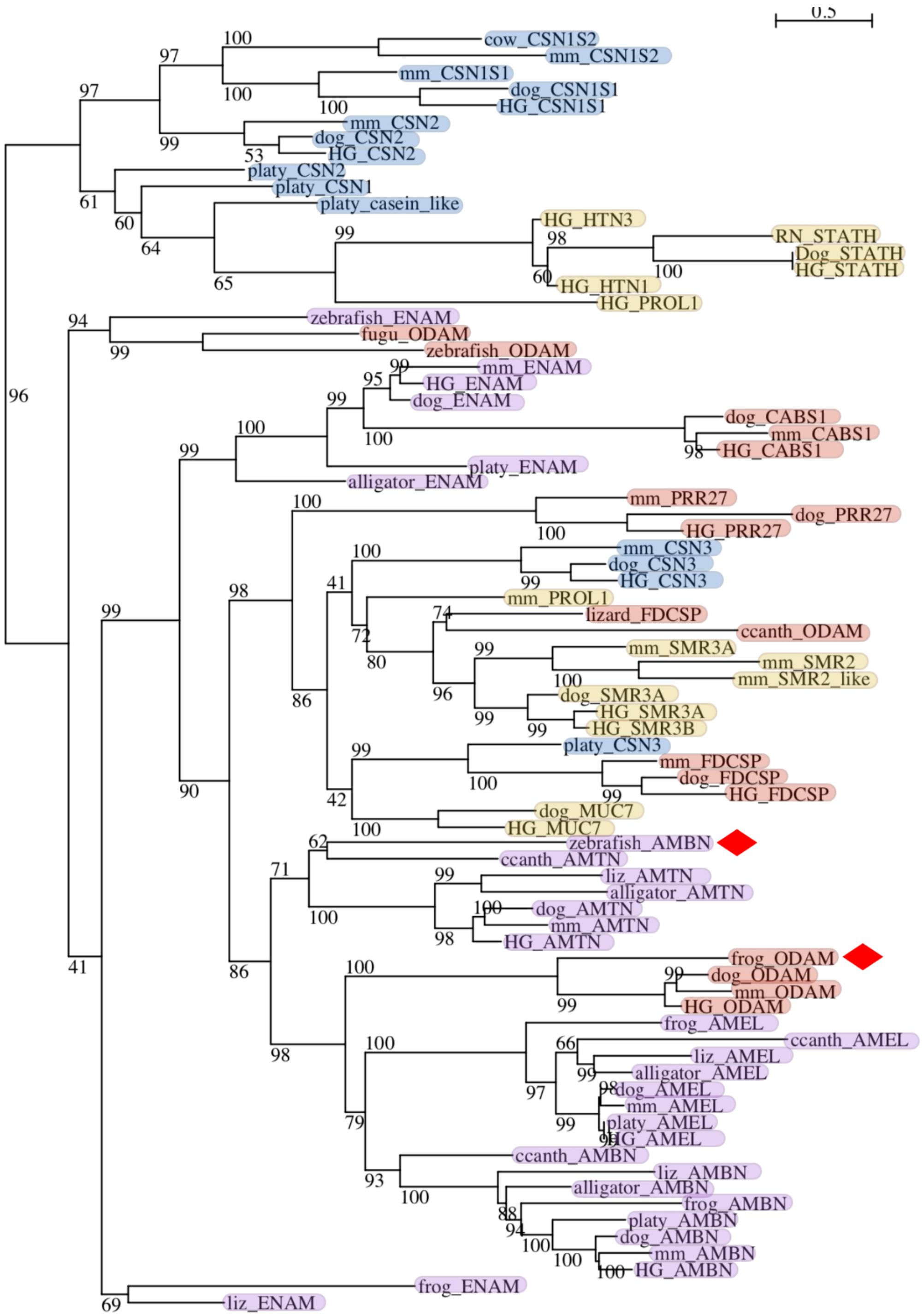


**Supplementary Figure 3: Gene tree of enamel-related SCPP genes**. Maximum likelihood tree generated using coding nucleotide sequences from major clades (mammals [human, mouse, dog, platypus], reptiles/amphibian [lizard, alligator, frog], and fishes [coelacanth, fugu, zebra fish]). Bootstrap support is shown at the nodes. Genes are labeled with a species abbreviation followed by the gene name in capital letters. They are highlighted based on predetermined categorizations of functional expression (see **Table 1).** Red diamonds show potential convergent evolution where nucleotide level variation is clustered in two different branches, while amino-acid sequences are similar and cluster in a single branch. Species name abbreviations are as follows: HG (human), mm (mouse), platy (platypus), liz (lizard),and ccanth (coelacanth).

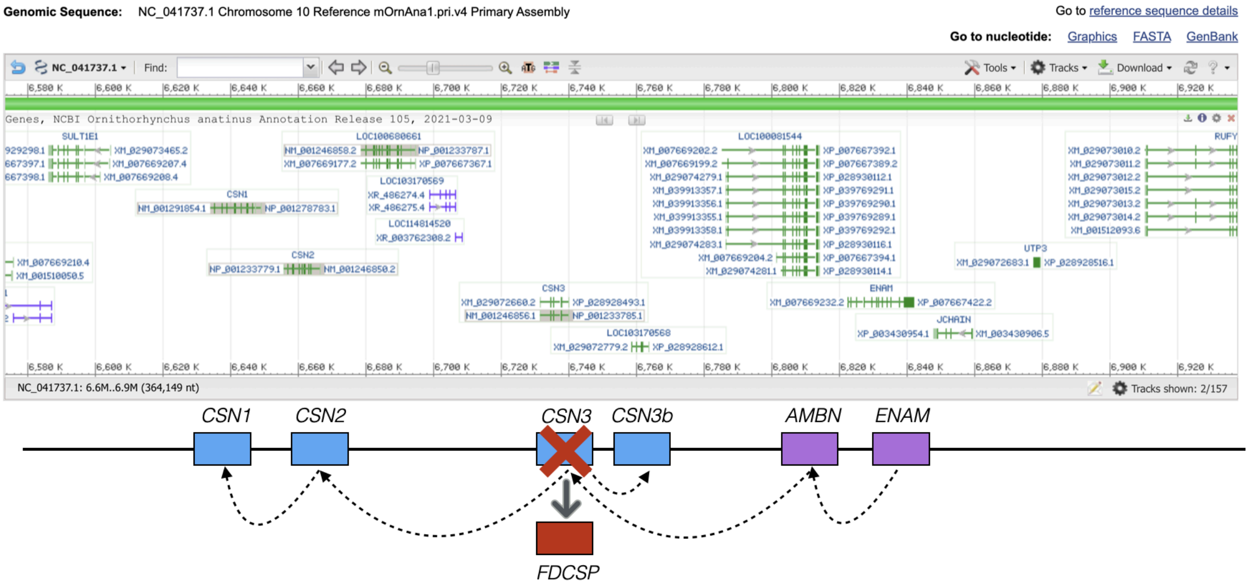


**Supplementary Figure 4: SCPP gene locus in platypus reveals a missannotation of *FDCSP* as *CSN3*.** A screenshot of NCBI genome browser, showing the platypus gene models and predictions for the SCPP gene locus. Genes without annotations are labeled as “LOC”. Below the screenshot is a simplified version of the models illustrating the likely ancestral gene duplication history of the SCPP genes in mammals, as proposed based on our findings. Genes are colored based on functional categorization: enamel-related genes (purple), milk-related casein genes (blue), and broadly-expressed genes (red). Arrows indicate the proposed paths of gene duplications.

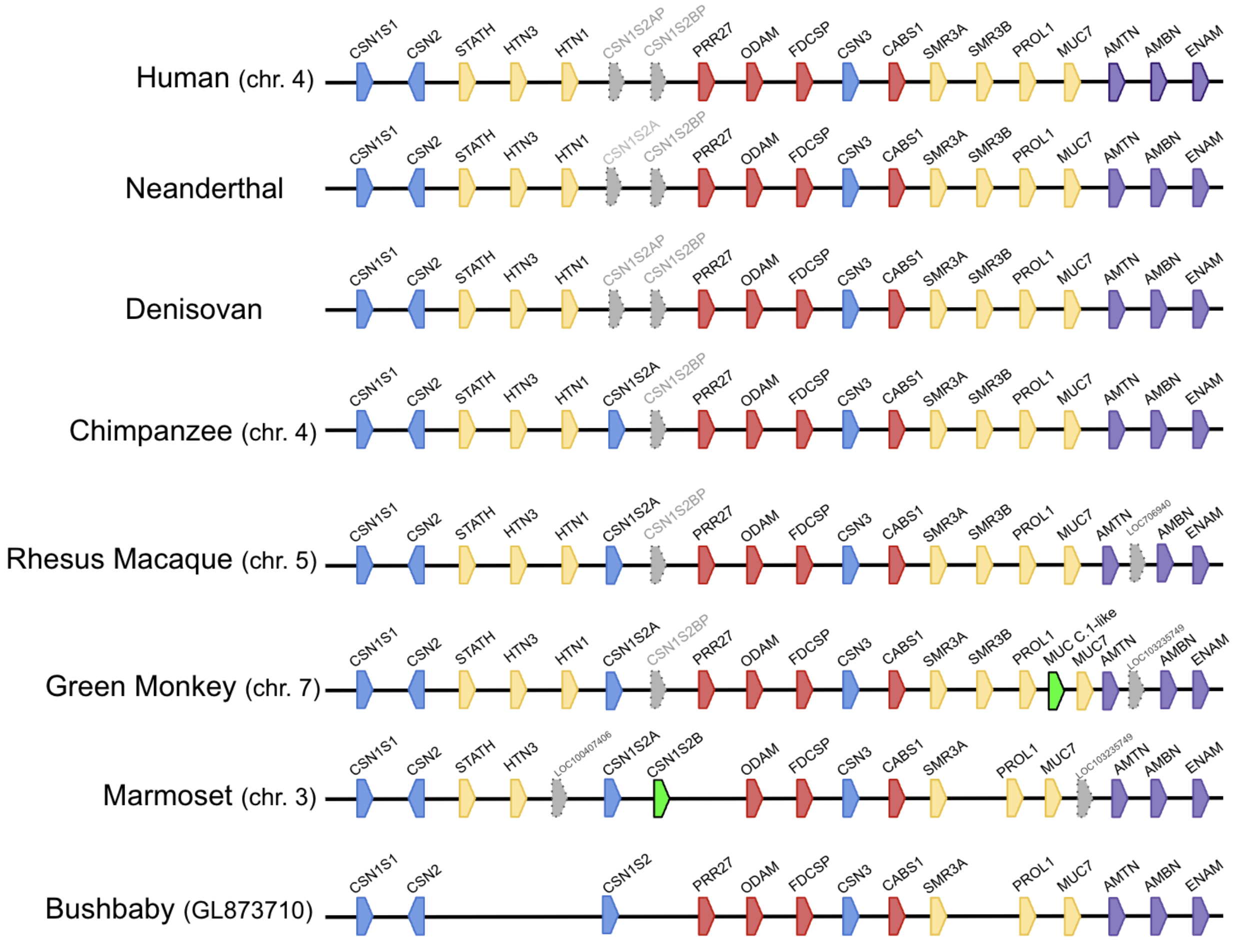


**Supplementary Figure 5: SCPP gene locus in primates shows the evolution of saliva-related genes, lineage-specific genes, and pseudogenization.** Genes are colored by expression patterns: enamel-related genes (purple), milk-related casein genes (blue), broadly expressed genes (red), and saliva-related genes (yellow). Pseudogenes are shown in grey with dotted outlines. Proposed lineage-specific gene duplications are colored green.

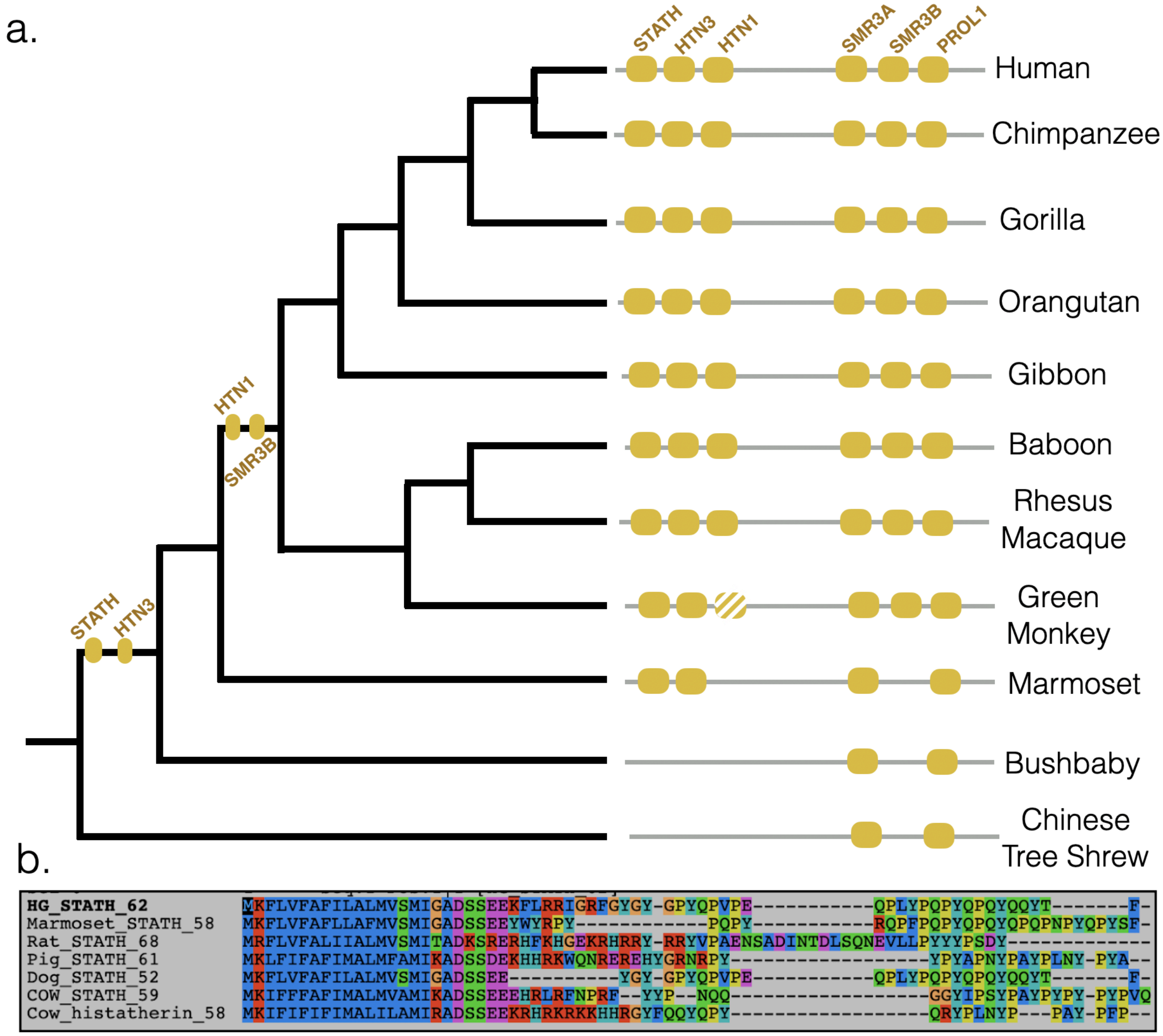


**Supplementary Figure 6: Sequential evolution of saliva-related genes in primates and alignment of STATH across species.** The top panel (a) shows the repertoire of saliva-related SCPP genes in a phylogeny of several primate species. Yellow boxes show the presence of each gene in each species, lower confidence presence of genes is indicated by striped boxes. The genes are also shown on the branch nodes, where they likely evolved through duplication from pre-existing genes within the SCPP locus. The lower panel (b) shows an alignment of amino acid sequences encoded by the *STATH* gene across several mammalian species, where it exhibits one main domain.

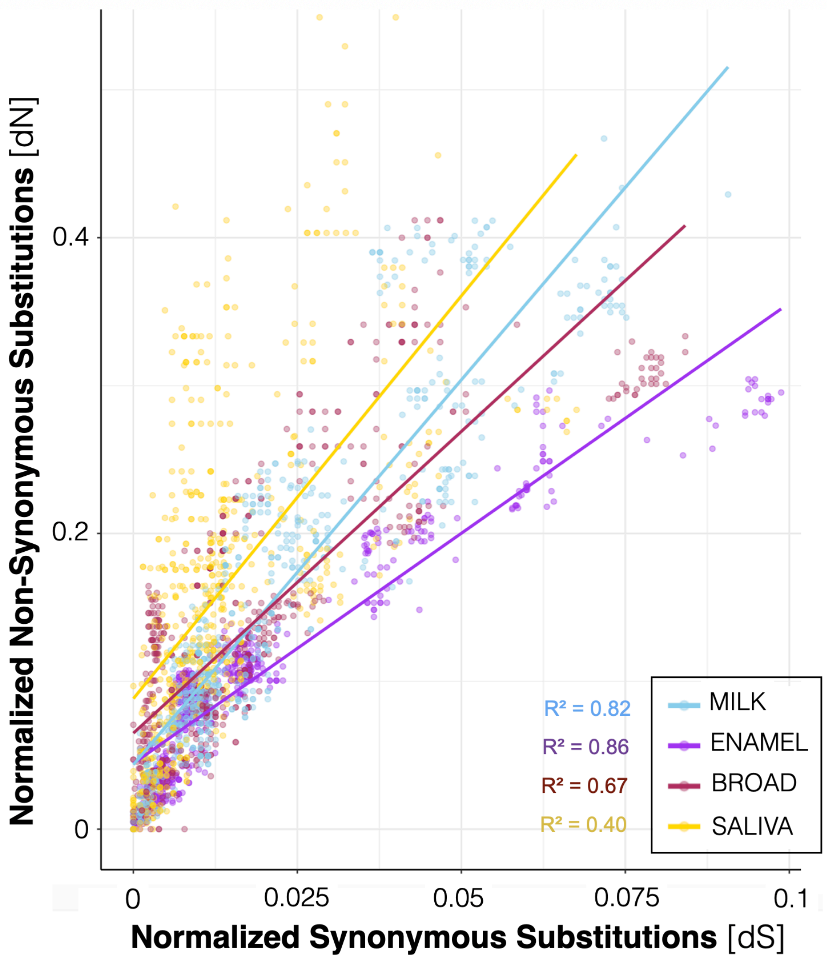


**Supplementary Figure 7: Evidence for selection on SCPP gene categories in primates.** Scatterplot of non-synonymous (dN,y-axis) v.s. synonymous nucleotide substitutions (dS, x-axis) in enamel-related genes (*ENAM*, *AMBN*, *AMTN*; purple), milk-related casein genes (*CSN1S1*, *CSN2*, *CSN3*; light blue), broadly-expressed genes (*PRR27*, *CABS1*, *ODAM*, *FDCSP;* red), and saliva-related genes(*MUC7*, *SMR3A&B*, *HTN1&3*, *STATH;* yellow). dN and dS values were normalized by total protein length and total transcript length, respectively. Linear regression lines are shown for each gene type with each functional category.
